# The Normative Modeling Framework for Computational Psychiatry

**DOI:** 10.1101/2021.08.08.455583

**Authors:** Saige Rutherford, Seyed Mostafa Kia, Thomas Wolfers, Charlotte Fraza, Mariam Zabihi, Richard Dinga, Pierre Berthet, Amanda Worker, Serena Verdi, Henricus G. Ruhe, Christian F. Beckmann, Andre F. Marquand

## Abstract

Normative modeling is an emerging and innovative framework for mapping individual differences at the level of a single subject or observation in relation to a reference model. It involves charting centiles of variation across a population in terms of mappings between biology and behavior which can then be used to make statistical inferences at the level of the individual. The fields of computational psychiatry and clinical neuroscience have been slow to transition away from patient versus “healthy” control analytic approaches, likely due to a lack of tools designed to properly model biological heterogeneity of mental disorders. Normative modeling provides a solution to address this issue and moves analysis away from case-control comparisons that rely on potentially noisy clinical labels. In this article, we define a standardized protocol to guide users through, from start to finish, normative modeling analysis using the Predictive Clinical Neuroscience toolkit (PCNtoolkit). We describe the input data selection process, provide intuition behind the various modeling choices, and conclude by demonstrating several examples of down-stream analyses the normative model results may facilitate, such as stratification of high-risk individuals, subtyping, and behavioral predictive modeling. The protocol takes approximately 1-3 hours to complete.

## Introduction

Clinical neuroscientists have recently acknowledged two realities that have disrupted the way research is conducted: first, that to understand individual differences it is necessary to move away from group average statistics ^1–7^ and, second, that the classical diagnostic labels of psychiatric disorders are not clearly represented in the underlying biology^8–11^. Initiatives such as RDoC^9,12,13^, HiTOP^10,14,15^, and ROAMER^16,17^ were established in response and seek to refine the nosology of mental disorders by mapping biobehavioral dimensions that cut across heterogeneous disorder categories. Despite this awareness and an increasing interest in quantifying individual differences, the field has still been slow to transition away from case-control comparisons that aim to contrast patient versus healthy control groups and assume that clinical groups are distinct and homogenous. A key barrier that has impeded progress is a lack of alternative analysis methods, designed to model variation across individuals, also known as heterogeneity^18^. Nearly all existing techniques for connecting the brain to behavior operate at the group-level and provide no path to individual-level inference^19–21^. Normative modeling is a framework for understanding differences at the level of a single subject or observation while mapping these differences in relation to a reference model (Figure 1). It involves charting centiles of variation across a population in terms of mappings between biology and behavior, which can then be used to make statistical inferences at the level of the individual, akin to the use of growth charts in pediatric medicine (Figure 1A). The practice of normative modeling in clinical neuroscience was developed to provide additional information beyond what can be learned from case-control modeling approaches (see ‘Development of the Protocol’ section below for further information). Case-control thinking assumes that the mean is representative of the population, when it may not be (e.g., if the clinical population is diffuse or comprised of multiple sub-populations). Therefore, normative modeling has become a leading tool for precision medicine research programs and has been used in many clinical contexts^22^ (see ‘Applications’ section below for further examples).

**Figure 1.**
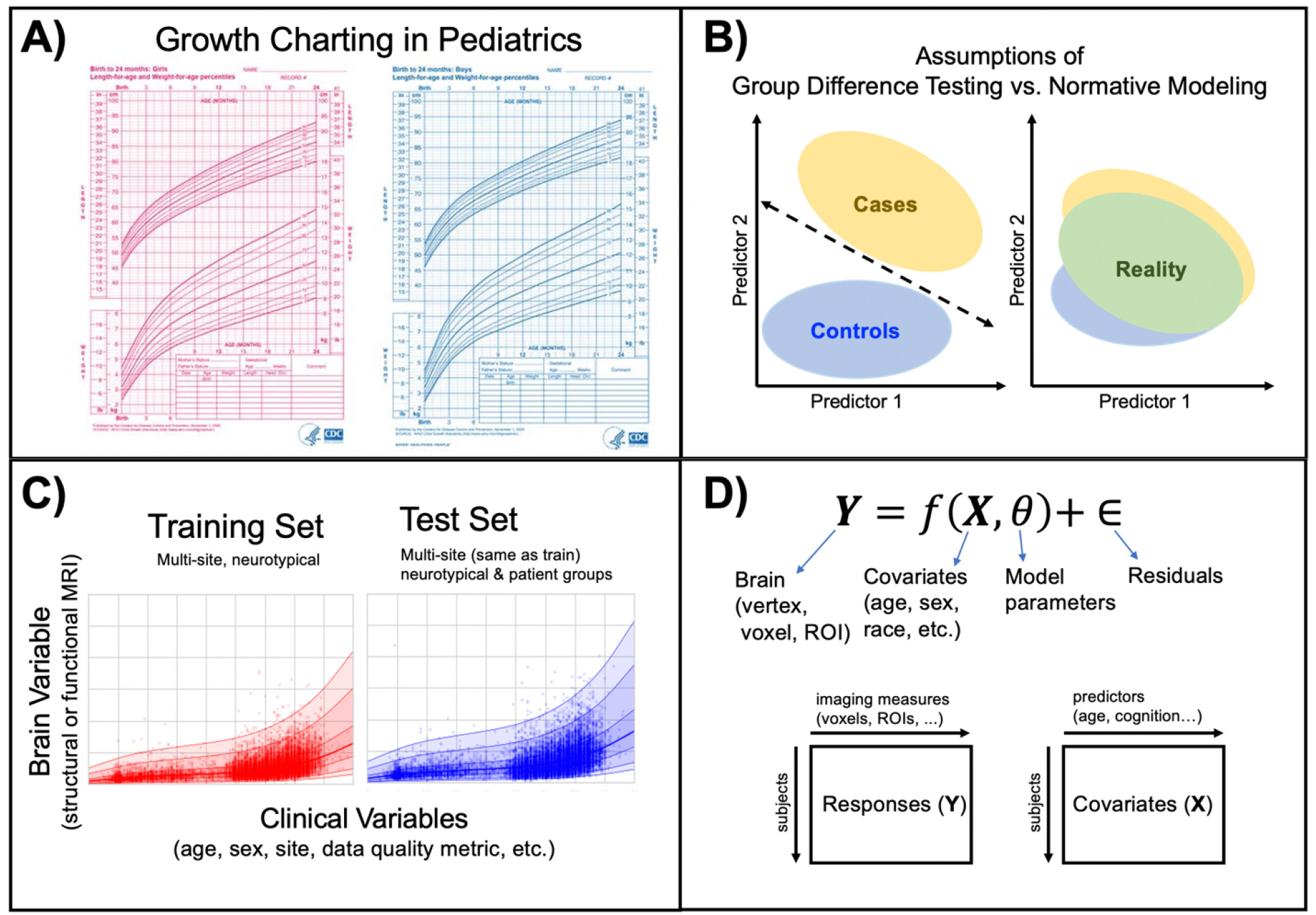
Conceptual Overview of Normative Modeling. **A)** Classical example of normative modeling: the use of height and weight growth charting in pediatrics. **B)** Case-control models (left) theoretically make assumptions that there is a boundary that can separate groups and that there is within-group homogeneity. In reality (right), there is nested variation across controls and patient groups and within-group heterogeneity, resulting in unclear separation boundaries. Normative modeling is well equipped to handle this reality. **C)** An example application of normative modeling in computational psychiatry using neuroimaging data. Mean cortical thickness (y-axis) is predicted from age (x-axis) using a training set consisting of multi-site structural MRI from neurotypical controls and a test set consisting of neurotypical controls and patient groups. Every dot indicates the deviation score for a single individual from normal development. **D)** Regression model equation and design matrix setup for the model shown in panel C.

Neuroscience has historically brought together scientists from diverse educations, for example, some from a clinical background and others having a mathematics background. The interdisciplinary nature introduces a challenge in bridging the gap between technical and clinical perspectives. This is a key challenge that aligns with the aims of the open-science movement and brain-hack community^23^, in other words, to distill the essential components of the analytic workflow into a consistent and widely applicable protocol. This helps to avoid ‘research debt’, i.e., a lack of ideas being digested^24^. This distiller mindset is crucial for confronting research debt and embracing paradigm shifts in thinking, such as moving from case-control comparisons to the normative modeling framework.

The purpose of this work is to distill the methods of normative modeling, an advanced analysis technique, into an actionable protocol that addresses these challenges in that it is accessible to researchers within the diverse field of clinical neuroscience. We distill the essential components of a normative modeling analysis and provide a demonstrative analysis from start to finish using the Predictive Clinical Neuroscience Toolkit <u>software</u>. We describe the input data selection process, give an overview of the various modeling choices, and conclude by demonstrating several examples of downstream analyses the normative model results may facilitate, such as stratification of high-risk individuals, subtyping, and behavioral predictive modeling.

## Development of the protocol

Normative modeling has a long history that relates to statistics and measurement theory and has many applications from medicine to economics to neuroscience. Familiar use cases of normative modeling include growth charting in pediatrics, neurocognitive tests, and interpreting graduate school test score percentiles (i.e., scoring 90^th^ percentile on the MCAT). The mathematical and computational development of normative modeling has been fine-tuned^25–28^ and currently exists as an open-source software python package, the Predictive Clinical Neuroscience toolkit (PCNtoolkit), which we focus on in this manuscript. This toolkit implements many commonly used algorithms for normative modelling and supports multiple industry standard data formats (e.g., NIFTI, CIFTI, text formats). Extensive documentation has been written to accompany this protocol and is available online through <u>read the docs</u>. This includes tutorials with sample data for all algorithm implementations, a glossary to help new users understand the jargon associated with the software, and a frequently asked questions page. An online <u>forum</u> for communicating questions, bugs, feature requests, *etc.* to the core team of PCNtoolkit developers is also available. We have developed these open-source resources to promote and encourage individual differences research in computational psychiatry using normative modeling.

## Applications and comparison with other methods

Normative modeling has been applied to many research questions in computational psychiatry and other fields, including in autism spectrum disorder^29–31^, attention deficit hyperactive disorder^32,33^, Alzheimer’s disease^34^, bipolar disorder, and schizophrenia^35–37^. Crucially, these applications have shown that normative modelling can detect individual differences both in the presence of strong case-control differences (observed in schizophrenia)^36^ and in their absence (observed in autism spectrum disorder)^30^. This highlights the value and complementary nature of understanding individual variation relative to group means. These applications have primarily focused on predicting regional structural or functional neuroimaging data (i.e., biological response variables) from phenotypic variables (i.e., clinically relevant covariates) such as age and sex. Age creates a natural, time-varying dimension for mapping normative trajectories and is well suited to applications in which deviations of an individual manifest from a typical trajectory of brain development or ageing. However, other phenotypes that have been used in neuroimaging predictive modeling studies such as general cognitive ability^38,39^, social cognition, or sustained attention^40,41^ are also attractive possibilities to use as covariates, thereby defining axes for observing deviation patterns. Normative modeling has also been used to learn mappings between reward sensitivity and reward related brain activity^42^.

It is important to emphasize that normative modeling is a general regression framework for mapping sources of heterogeneity, refocusing attention on individual predictions rather than group means (e.g., diagnostic labels), and detecting individuals who deviate from the norm. Therefore, it is not limited to a specific algorithm or mathematical model, although we recommend certain algorithms based on the research question and available input data. The algorithms in the PCNtoolkit tend to favor Bayesian over frequentist statistics, as there are certain features of Bayesian approaches that facilitate better normative modeling estimation. For example, having a posterior distribution over the parameters help to better separate different sources of uncertainty, e.g., separating variation (‘aleatoric uncertainty’ – cannot be reduced by adding more data) from modeling (or ‘epistemic’) uncertainty which can be reduced by adding more data. These different use cases of normative modeling (algorithm selection, predicting brain from behavior or behavior from the brain) are explained in-depth in the ‘Experimental Design’ section, below.

There is a long history of using regression methods to learn mappings between brain and behavior^43,44^. Principle Component Regression (PCR)^20,45,46^, Connectome predictive modeling (CPM)^19,47^, and canonical correlation analysis (CCA)^48,49^ have become mainstream methods for linking brain and behavior. These methods have demonstrated the feasibility of brain-behavior mapping and laid the foundation for individual differences research to thrive. While these approaches have generated much curiosity and excitement, they are limited in their ability to provide inference at the level of the individual, providing only point estimates (i.e., without associated centiles of variation). Most papers using these tools only report the mean predictive model performance, collapsing information across hundreds or thousands of people into a single number (e.g. model accuracy or regression performance)^46,47,50,51^. The normative modeling framework takes these ideas a step further to quantify and describe how individuals differ statistically, with respect to an expected pattern. In this way, normative modelling breaks the symmetry inherent in the case control paradigm. In more detail, PCR and CPM differ from normative modeling in terms of how the prediction model is formulated. PCR and CPM setup the regression model such that, Y, a n_subjects x 1 vector (i.e., age or fluid intelligence), is predicted from X, a matrix with n_subjects x n_brain dimensions, where n_brain is typically a reduced feature space selected via a regularization step. This setup makes interpretating which brain features are related to the behavior very challenging. Studies using PCR or CPM attempt to interpret the brain feature weights, but as these methods typically use fMRI connectomic data, consisting of connections and nodes, interpretation often yields a complex whole-brain visualization that is not very informative. The individual-level output of these models is a single point estimate, a predicted behavior score for each subject. These individual point estimates are then summarized by correlating the predicted and true behavior scores, reporting explained variance (R^2^), and calculating accuracy (mean squared error). Compared to PCR and CPM, normative modeling inverts the regression setup around to predict brain region Y, a n_subjects x 1 vector from X, a matrix with n_subjects x n_covariates (i.e., age, sex, fluid intelligence, site, data quality metric). There is a separate regression model for each brain region. The individual-level outputs of normative modeling are the predicted brain score, the predictive variance (separated into modeling and noise components), a deviation score (Z-score, how much each subject deviate from the normative range). The overall performance is evaluated by correlating predicted and true values, calculating explained variance, standardized mean squared error, and mean standardized log loss. In contrast, CCA estimates a doubly multivariate relationship in that both X and Y are matrices (X is n_subjects x n_brain matrix and Y is n_subjects x n_behavior). Whilst CCA is well suited to detecting that a mapping exists, this still leads to difficult interpretation of feature importance and, moreover, CCA is highly prone to overfitting and requires careful assessment of out of sample metrics with respect to an appropriate null distribution, which is not always done in practice. Like PCR and CPM, CCA also does not provide individual measure of uncertainty or deviation scores.

Case-control inference (e.g., mass univariate group t-testing and classification of patient vs. control) examples are perhaps the most interesting comparison to the normative modeling framework. Case-control methods typically require there to be a homogeneous within-group spatial signature and their success relies on obtaining statistical significance (p-value < 0.05). We clarify this point with an example of the assumptions of case-control inference. To detect a group difference in amygdala activation between a control group and a group of individuals with post-traumatic stress disorder during a fMRI task, all individuals in the control group need a similar value of amygdala activation and all individuals in the PTSD group need a similar value of amygdala activation. Then, the mean amygdala activation signal of the control group must be statistically different from the mean amygdala activation signal of the PTSD group after stringent multiple comparison testing correction. These assumptions ignore the fact that different biological processes (i.e., some people have increased activation and others decreased) can lead to similar external behavior. Normative models reveal a different side of the data -- that the classical diagnostic labels of psychiatric disorders are not clearly represented in the underlying biology, meaning patient groups are not well defined by a unifying neurosignature -- and provide clear evidence for the limitations of case-control paradigms. Brain age models are also in the same family as normative models but generally have a narrower focus on interpreting accelerated/decelerated aging^52,53^ or improving prediction accuracy^54^. Brain age models only allow for interpreting centiles of variation in terms of age, which is limited and does not have a clear interpretation in terms of biological variation across individuals.

“All models are wrong, but some are useful” -- George E.P. Box.

There is not one ‘best’ modeling approach and many of the methods presented in this section can be complimentary in that they investigate different questions. Before embarking on a computational modeling journey, it is always important to ask questions such as: What are the assumptions made by this model? What type of inference do you want to make (group-level, individual-level)? What aspect of the predictive model is most important (accuracy, quantifying uncertainty, statistical significance)? Allow your research question to guide the answers and model selection.

## Expertise needed to implement the protocol

We aimed to make this protocol user-friendly to the diverse community of neuroscience, including those with a non-technical background. The fundamental objective of this protocol is to learn how to implement the normative modeling framework via the PCNtoolkit software without being an expert in statistics and machine learning. You will be given enough knowledge to set up training and test sets, understand what data should be going into the model, interpret results, and make inferences based on the results. Prerequisites of this protocol are basic familiarity with the Python programming language and a computer with a stable internet connection. Complete code, example data, and extensive documentation accompany this protocol; thus, writing code from scratch is unnecessary. Of course, it is our intention for readers to be inspired by this protocol and to use the normative modeling framework in more ways than presented here. If you wish to use the framework presented in this protocol beyond the provided code, familiarity with the Linux command line, bash scripting, setting up virtual environments, and submitting jobs to high-performance clusters would also be helpful.

## Limitations

### Big data requires automated QC

As datasets grow to meet the requirements of becoming population-level or big data, there is typically a need to rely on automated quality control metrics^55^. This means there is potential to unintentionally include poor quality data, which could, in turn, affect the results. The training and test dataset used in this protocol has been manually quality checked by visualizing every subject’s raw T1w volume with their corresponding Freesurfer brain-mask as an overlay using an online (JavaScript-based) image viewer. Quality checking code and further instructions for use is made available on GitHub. These images were inspected for obvious quality issues, such as excess field-of-view cut-off, motion artifacts, or signal drop-out. Subjects that were flagged as having quality issues were excluded from the sample. Users should consider manually quality checking their own data if they wish to add on additional samples to the dataset.

### Multi-site confounds and data availability

Pooling data from multiple sites is often a necessary step to create diverse datasets and reach sufficient sample sizes for machine learning analyses. When combining data from different studies, several challenges arise. First, there are often different MRI scanners at each site that also have different acquisition parameters. These MRI hardware and software divergences give rise to substantial nuisance variance that must be properly accounted for when modeling the data. Second, there may be sampling differences, for example due to different inclusion criteria and definitions of diagnostic labels at each site. For example, if one site uses the Structured Clinical Interview for DSM-5 (SCID-5) administered by a trained mental health professional who is familiar with the DSM-5 diagnostic criteria, while another site relies on self-report questionnaire data to define clinical labels. This increases the heterogeneity within the clinical groups (e.g., by mixing inclusion criteria across cohorts) and could also add noise to the diagnostic labels (e.g., if diagnostic assessments have different reliability across studies). This is important to consider if clinical labels are used to separate data into training and testing sets (i.e., controls only in the training set) and when comparing the outputs of normative modeling across patient groups from different sites.

Furthermore, there is likely to be dissimilarities in the available demographic, cognitive, and clinical questionnaire data across sites as well which needs to be considered when deciding which studies to include and which covariates to use in modeling. If the goal is to share the model, allowing transfer to new samples, using unique covariates that are specific to your sample (and not commonly collected) will hinder the ability of others to use your model on their own data. There is a careful balance that should be considered regarding the benefits gained from a new site joining the sample versus the site related nuisance variance that accompanies the addition of new sites.

## Overview of the procedure

### Experimental design

There are many choices and considerations that should be carefully planned before embarking on a normative modeling analysis – the decision points can be grouped into the following stages: data selection, data preparation, algorithm/modeling, and evaluation/interpretation. These stages, and the corresponding step numbers of the procedure are summarized in Figure 2. There are additional resources and support for running normative modeling analysis that are summarized in Figure 3.

**Figure 2.**
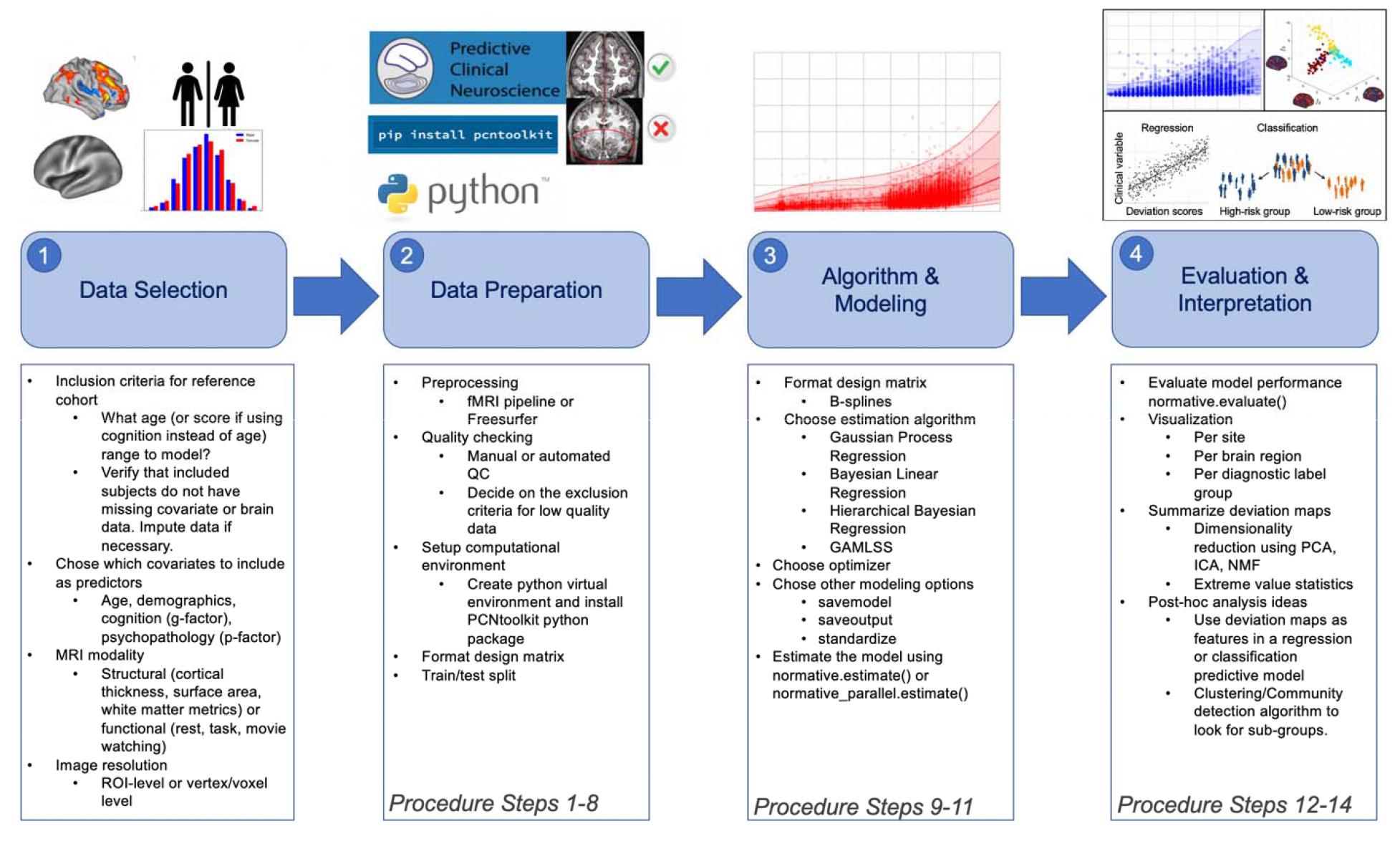
Practical Overview of Normative Modeling Framework. The workflow consists of four stages: data selection, data preparation, algorithm & modeling, and evaluation & interpretation, which are visualized by the numbered shaded blue boxes. The steps involved at each of these stages are summarized in the box below and highlighted in the images above.

**Figure 3.**
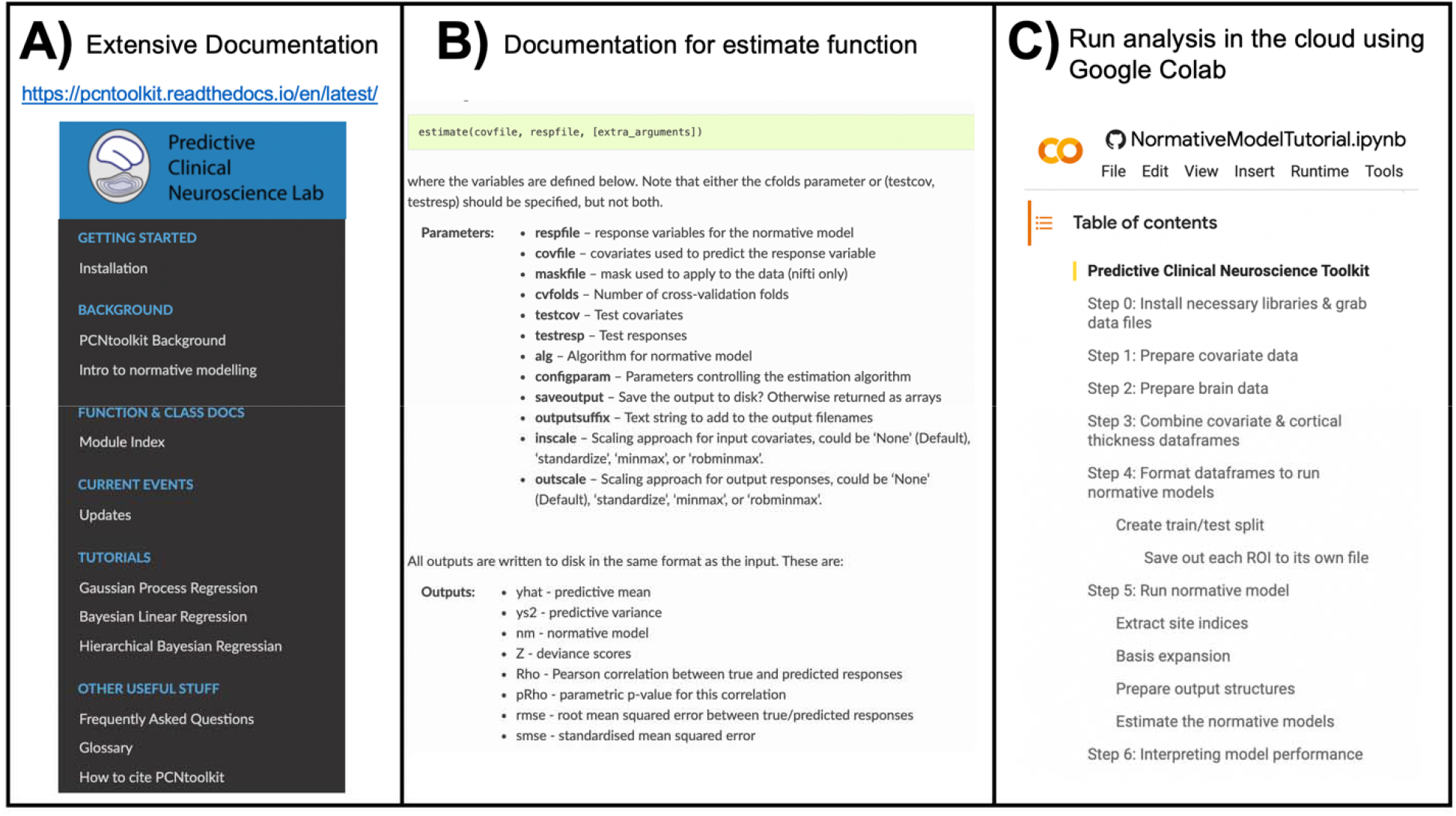
Overview of Resources for Running a Normative Modeling Analysis. **A)** Detailed documentation, including installation instructions, input/output descriptions of all classes and functions implemented in the python package, tutorials for all algorithms, frequently asked questions, a glossary explaining acronyms and other jargon, references to existing normative modeling literature, and a citation guide, is available <u>online</u>. **B)** Example of the documentation showing the required input, expected output of the main function used in the pcntoolkit software, the estimate function. **C)** All of the code and data used in this protocol is available to run in the cloud via Google Colab. Additional tutorials (shown under the tutorials header in panel A) are also available to run in Google Colab.

### Data selection – reference cohort inclusion criteria

Creating the training dataset that will serve as the “normative” reference cohort is the first important decision. Ideally, the training dataset will be a large and representative sample, and the included subjects should not be missing vital demographic (age, sex) or biological (neuroimaging) data. However, data imputation may be used if necessary but should be used cautiously. In most research studies, data are missing not at random, and we interpret more than just mean effects. In this case, mean imputation may bias results and other forms of imputation should be considered^56,57^. It is important that the reference cohort provides good coverage (complementary covariates) of the test set (e.g., clinical) population.

The sample size of the reference cohort (training set) is important to consider in normative modeling, although we emphasize that the focus is different to classical power calculations, which target a fundamentally different question (i.e., determining a required sample size to detect a group level comparison of a specified effect size at a given significance level). In contrast, in normative modeling, the focus is usually on quantifying deviations from a reference model at the individual level. In this context, the size of the reference cohort primarily influences the test set deviation scores by influencing the accuracy and precision with which the target phenotype (i.e., response variable) can be predicted. As the sample sizes increase, the predictive intervals will shrink, which results in an increased sensitivity to detect individual differences. However, there is not a specific cutoff that represents an ideal sample size, and we emphasize that context is key. For instance, you could build a clinical normative model for a sample of individuals with major depression disorder (MDD) for the purpose of stratification or detecting subgroups (individuals who have recurrent episodes or individuals who do not respond to medication). In this case, the reference cohort might consist of individuals that have experienced single MDD episodes and those that have responded well to medication. The sample size for this normative modeling research question would likely be relatively small due to data availability (e.g., clinical datasets typically have stricter data sharing requirements). The main takeaway from this MDD example is that sample size is highly dependent on the research question which in turn guides the inclusion criteria for the reference cohort you want to measure deviations from. If you are modeling “healthy” lifespan populations, the sample size will likely be large (on the order of thousands) because of the plethora of publicly shared data that can be leveraged. On the other hand, if you want to model a specific clinical population or a specific functional task, the sample size will be smaller due to availability of data. A smaller dataset that properly addresses the research question at hand is completely acceptable.

### Data selection - covariate selection

The next choice should be regarding which covariates to include. One of the main criteria to include a covariate is the relevance to the posed research question. In normative modeling, usually we are interested in studying the deviations from the norm of the population, in other words, we are more interested in residuals. Thus, when we include a covariate in the design matrix for estimating the normative model, we are mainly interested in removing its effect from the residuals (thus deviations) than investigating its effect on the neuroimaging variable. Normative modeling is a tool to study unknowns (that are encoded in the deviations). To do so, we need to first account for known variation in the data by regressing them out of the data (thus we include the knowns in the covariates), and then we interpret the residual variation in the deviation scores. For example, if you want to know the effect of smoking on the ventral striatum, that is not confounded by other substance use, you should include substance use variables (e.g., drinks per week, etc.) in the covariate matrix, estimate the normative model, and then correlate the ventral striatum deviation score (that has the effect of drinking removed from it) with smoking frequency. When pooling data from multiple sites, the available measures across sites may influence the selection of covariates because ideally, the variables should be consistent across sites. For example, you should not use different versions of a cognitive test, as they could test for different dimensions of general cognitive ability. For neurodevelopmental or lifespan model, the suggested minimum covariates to include are age, sex, site (using random- or fixed-effects), and optionally a metric of data quality (i.e., mean framewise displacement or Freesurfer Euler number). Modeling site is very important; however, an exhaustive explanation is outside the scope of this protocol but see ^20,21^ for an in-depth account of modeling site variation. Diagnostic labels could also be included as covariates to utilize the variance explained by these labels without constraining the mapping to only reflect case-control differences. Furthermore, additional biological covariates could also be included, for example blood biomarkers, or structural brain measures if predicting functional brain measures. Additional or alternative covariates may include other demographics (race, ethnicity, gender, education level, marital status, household income) and cognitive variables.

### Data Selection – MRI modality and spatial resolution of brain data

Next, it is necessary to decide on the modality of brain imaging to model. In this protocol, we use cortical thickness and subcortical volume measurements from structural MRI (T1-weighted) images. However, other modalities such as resting-state and task-based functional MRI or diffusion weighted MRI could also be selected in this step (data for these modalities is not provided with this protocol). The resolution of brain data is important to consider while keeping in mind the increasing computational complexity with modeling smaller units. Vertex or voxel-level modeling of brain data provides high-resolution deviation maps. Still, region of interest (ROI) level modeling may allow for easier interpretation/visualization of the output deviation maps and will have a lower penalty in multiple comparison correction (if doing post-hoc analysis) on the deviation maps. The PCNtoolkit can run models in parallel to speed up computation time; however, there is still a univariate nature, meaning a separate model is fit for each brain region. This univariate approach does not address the spatial autocorrelation^58–65^ or functional heterogeneity (functional mis-registration) present in (f)MRI data^66^. Spatial autocorrelation refers to the complex spatial correlation patterns present in MRI data. Nearby regions are often more correlated than distant regions, thus they are not statistically independent. Spatial correlations are difficult to model due to their heterogeneity, complexity and high dimensionality with a limited sample size. Techniques such as Markov Random Fields^62,63^, network/graph theory (topology)^58,59,64,65^, and spatial Bayesian latent factor methods^61,67^ have been applied to address the problem of spatial autocorrelation in raw or preprocessed MRI data. Progress in addressing spatial autocorrelation in the context of normative modeling has also been made in which Kronecker algebra and low rank approximations are used to build multivariate normative models^68,69^. In the context of normative modeling, we recommend paying extra attention to image registration in order to properly model the functional regions, as the spatial overlap of regions across individuals is not guaranteed with functional areas. In addition to taking extra measures to align the fMRI data, rather than modeling single voxels or parcels (as is often done in structural MRI), it may be beneficial to model brain networks as these features better capture the spatial patterns of functional units.

### Data preparation – Preprocessing and quality checking

Example data has been curated and shared for the purposes of this protocol. As mentioned in the data selection – MRI modality section above, we use structural MRI and have run Freesurfer to extract cortical thickness and subcortical volume measures. If using other data than the provided protocol data (i.e., your own data) then you will be to preprocess it accordingly and quality check the data to ensure only high-quality data is included. If you are new to working with MRI data, we recommend <u>Andy’s Brain Book</u>^70^ that includes videos and code tutorials for most neuroimaging software (i.e., Freesurfer, FSL, SPM).

### Data preparation – Setup computational environment

In this stage you will create a python virtual environment and install the required python packages. Then you will clone the GitHub repository which contains all the code and data required to follow along with the procedure section. You can run the entire protocol in the cloud using Google Colab or chose to run the code on your own computer or server.

### Data preparation – Format design matrix (site effects)

It is rare for a single scanning site to acquire large enough samples that are an accurate representation of the general population. Therefore, it is common to pool data obtained across multiple MRI centers. Some projects, such as the ABCD study^71^, have begun to harmonize scanning protocols because multi-site pooling was planned prior to data collection. In contrast, other projects, such as ENIGMA^72^, combine data post-collection and not have harmonized scanning sessions prior to data collection. If possible, to eliminate additional sources of variance, multi-site pooled data should be preprocessed using identical pipelines and software versions. However, due to data sharing restrictions and privacy concerns regarding health data, raw data may be unavailable, making pre or post data collection harmonization efforts impossible. Data harmonization techniques, such as COMBAT^73–76^, aim to remove site-related variance from the data as a preprocessing step before further analyses are run. There are some issues with harmonization, principally that all sources of variance that are correlated with the batch-effects (i.e., site-related variance) are removed which can unintentionally remove important, unknown, clinically relevant variance from the data. COMBAT also requires that the user have access to all the data when harmonizing which may have implications for data privacy. We therefore do not recommend users focus on data harmonization techniques when preparing their data sets for normative modeling. Hierarchical Bayesian Regression (HBR)^27,28^ implemented in the PCNtoolkit has been thoroughly developed and tested to address these challenges when using multi-site data in normative modeling. HBR estimates site-specific mean effects and variations in the normative model estimation stage using a Bayesian hierarchical model, which produces site-agnostic deviation scores (z-statistics). This distinction between harmonization techniques (i.e., COMBAT) and HBR-normative modeling is very important when using deviation scores as features in subsequent interpretation analyses, as harmonization has been shown to overexaggerate confidence in downstream analyses^77^.

### Data preparation - Train-test split

Whilst there are no hard rules for selecting the relative proportion of training and test data, some general guidelines that may help this decision can be considered. On the one hand, it is important to ensure the training set be sufficiently large to model the target phenotype with sufficient accuracy and precision. On the other hand, ensuring the test set is not too small is also important to provide sufficient sensitivity to detect downstream differences (which may depend on the expected frequency of clinically relevant deviations in the test set). In practice a 70% train, 30% test or 80% train, 20% test split often provides a reasonable balance between these competing objectives, but in certain applications it may be necessary to deviate from these recommendations. The main purpose of the train-test split is to establish out of sample generalizability and whether there is over (or under) fitting occurring. More important than the exact ratio of the train/test split, we believe it is critical to focus on preserving the sample characteristics across the train/test split. For example, it would not be sensible to model age ranges of childhood and adolescence in the reference cohort and have the test cohort consist of late adulthood ages. This scenario would detect high deviations in this test set due to not properly modeling the target population. If you want to investigate the hypothesis that a certain clinical group (e.g., individuals with a psychosis diagnosis) have more extreme deviation patterns than a control group (individuals with no psychiatric diagnosis), you need to verify that it is because they are patients not because they are in the test set. In order to verify this, it is important to also include some controls (from the same imaging site as the patients) in the test set. In other words, you cannot separate site variation from diagnostic variation if you do not have control reference data.

The train-test ratio decision naturally relates to the sample size requirements of the reference cohort mentioned in the data selection – reference cohort inclusion criteria section above, and the same consideration of the context needs to be taken when creating the train-test split. Does this split align with the research question being asked? More specifically, does the training set adequately match the reference (“normative”) cohort and does the testing set represent the target cohort in which deviations (from the reference cohort) will be interpreted? We discourage cross validation, or iteratively resampling of the data set into train and test sets, unless the dataset is very small, and if it is used then practitioners should be aware of the problems it introduces. Ideally, the train/test split of the dataset will only be done once. While cross validation is useful for testing stability and sensitivity of models to perturbations, it also leads to having multiple models which are not easy to combine and interpret and it induces dependence between folds which violates most parametric statistical tests^78^.

### Algorithm & Modeling - algorithm selection

After the data have been carefully chosen and curated, it is time to move onto the normative modeling implementation. There are several algorithms for implementing a normative model including Gaussian process regression^79^, Bayesian linear regression^25,67^, hierarchical Bayesian regression^27,28^, generalized additive models of location, scale, and shape^26^, neural processes^68^, random feature approximation^80^, quantile regression^81^ and many of these are implemented in the PCNtoolkit software package (Table 1). The algorithms have different properties depending on their ability to model non-linear effects, scaling to large data sets (in terms of computation time), handling of random or fixed effects (e.g., to model site effects), their ability to model heteroscedastic or non-Gaussian noise distributions and their suitability for use in a federated or decentralized learning environment. An overview of these algorithm implementations is covered in Table 1. Gaussian process regression (GPR) was widely used in the beginning phases of normative modeling, which can flexibly model non-linear effects but does not computationally scale well when the training data increases (i.e., beyond a few thousand data points). In this work, we focus on Bayesian linear regression (BLR), which is highly scalable (fast compute time with large samples) and flexible (can be transferred to new sites not included in the training sample and can be combined with likelihood warping to model non-Gaussian effects). Hierarchical Bayesian regression (HBR) is another appealing choice as it has been used to better address multi-site datasets and allows for transfer learning (e.g., prediction for unseen sites) and can be estimated in a federated learning framework which is useful if there are privacy concerns and/or sharing restrictions meaning data cannot easily be pooled at a single computing site.

**Table 1.**
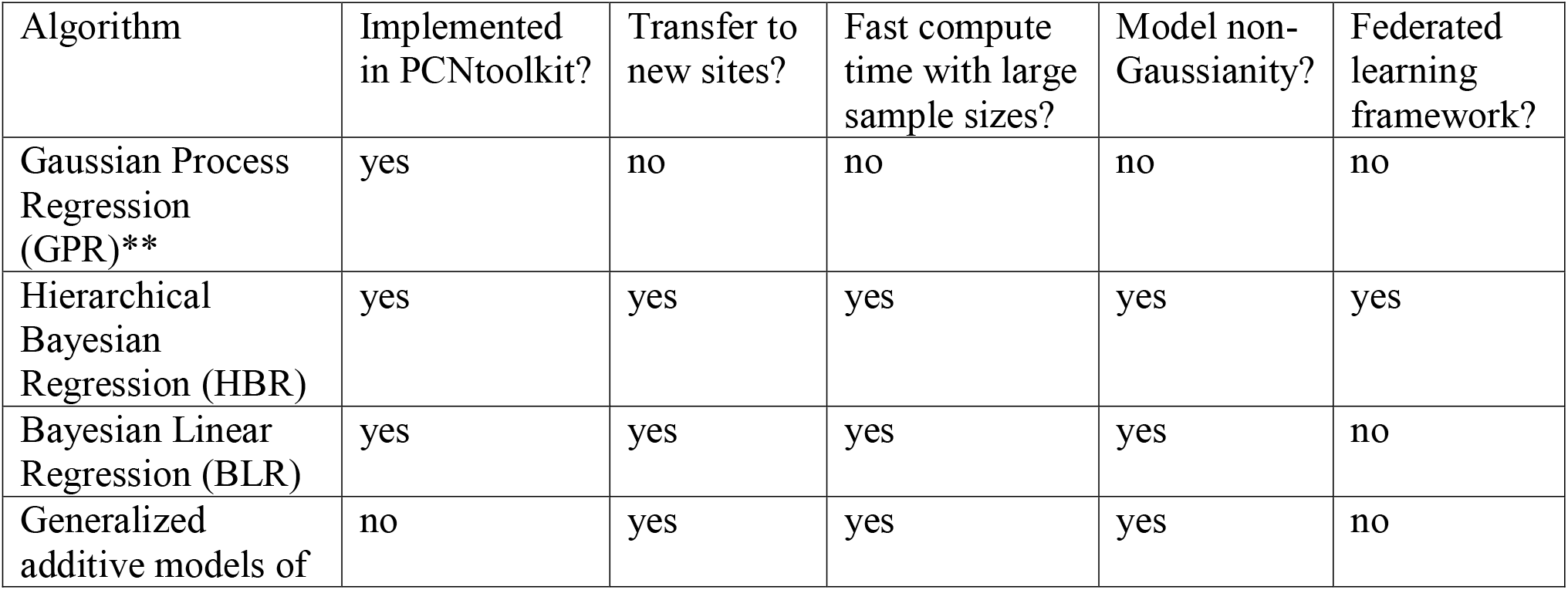
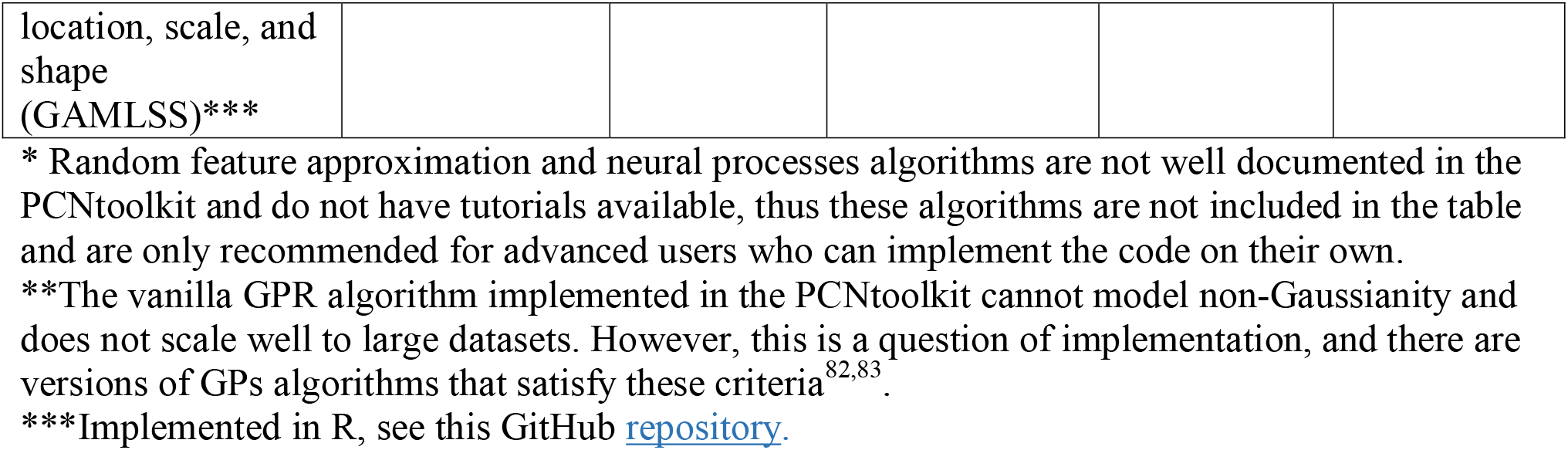
PCNtoolkit Normative Modeling Algorithm Overview*.

The remaining steps including estimating the normative model, evaluation the model performance, interpretating the model fit, and ideas for post-hoc analysis of the normative modeling outputs are covered in more detail in the protocol section.

## Materials

### Equipment

- Computing infrastructure: a Linux computer or HPC (SLURM or Torque) with enough space to store the imaging data of the train and test set.

- If a Linux computer or server is unavailable, this protocol can also be run in Google Colab (for free). If using Google Colab, only a computer with an internet connection and modern internet browser (e.g., Chrome or Firefox) installed is necessary.
- Python installation (https://www.python.org/downloads/).

- Recommended: Anaconda or virtual environment to manage the required python packages (https://www.anaconda.com/ or https://virtualenv.pypa.io/en/latest/).
- PCNtoolkit python package version 0.20 (and dependencies) installed via pip (https://pcntoolkit.readthedocs.io/en/latest/pages/installation.html).
- Covariates and response variables. Examples of these are provided with this protocol.

- Demographic and behavioral data used as predictor variables

- Age, sex/gender, site/scanner ID, race/ethnicity, cognition, data quality metric (Euler number if structural, mean framewise displacement if functional)
- Biological data to be modeled. An example structural MRI dataset is provided with this protocol.

- Structural MRI: cortical thickness, surface area, subcortical volume
- Functional MRI: parcellated task activation maps, resting-state networks

## Procedure

The data selection stage (Figure 2, panel 1) does not require code, as it is more of a research question formulation stage (i.e., choosing inclusion criteria and what type of imaging modality to model). Data preprocessing (running Freesurfer) and quality checking have also already been performed, and code for running Freesurfer or other preprocessing is not included in this protocol. Thus, for this protocol, the procedure begins at the *data preparation – setting up computational environment stage*. See the experimental design section for guidance on the data selection stage and preprocessing if using different data than what is provided with the protocol.

### Data Preparation: Prepare computational environment

#### Timing: 1-3 minutes

1. Begin by cloning the GitHub repository using the following commands. This repository contains the necessary code and example data. Then install the python packages using pip and import them into the python environment (either Google Colab or using a local python installation on your computer), as follows:

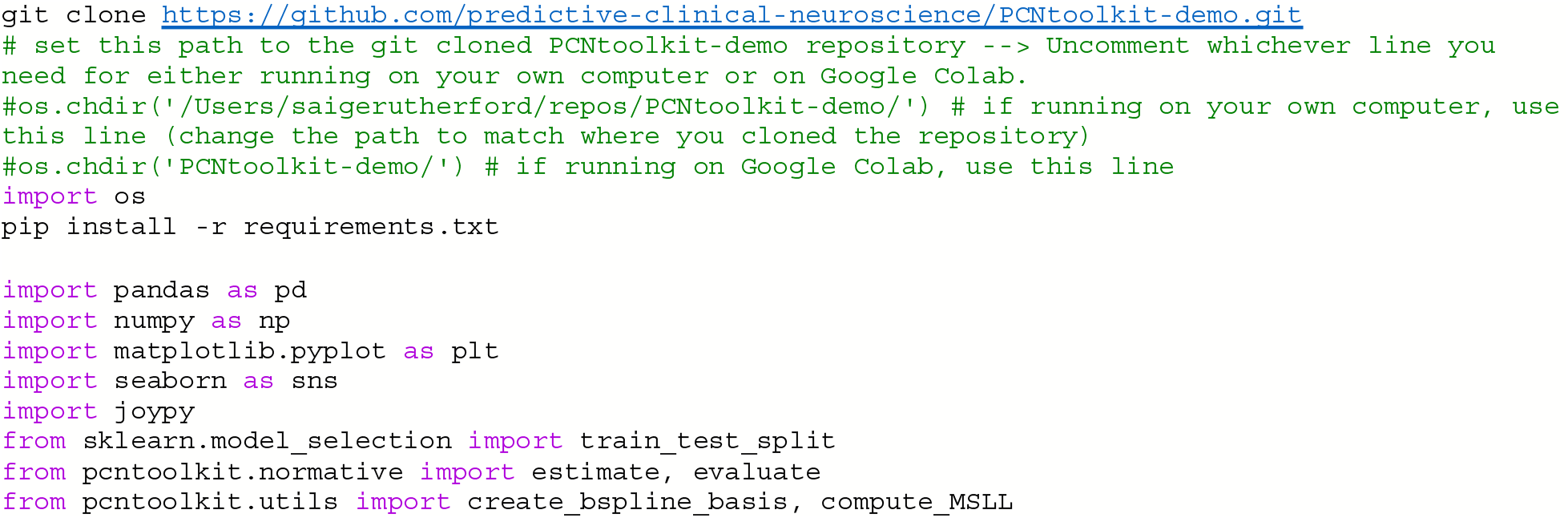

### Data Preparation: Prepare covariate data

#### Timing: 5-8 minutes

2. The data set (downloaded in Step 1) includes a multi-site dataset from the <u>Human</u> <u>Connectome Project Young Adult study</u>, <u>CAMCAN</u>, and <u>IXI</u>. It is also possible to use different datasets (i.e., your own data or additional public datasets) in this step. If using your own data here, it is recommended to load the example data to view the column names in order to match your data to this format. Read in the data files using pandas, then merge the covariate (age & sex) data from each site into a single data frame (named cov). The columns of this covariate data frame represent the predictor variables. Additional columns may be added here, depending on the research question.

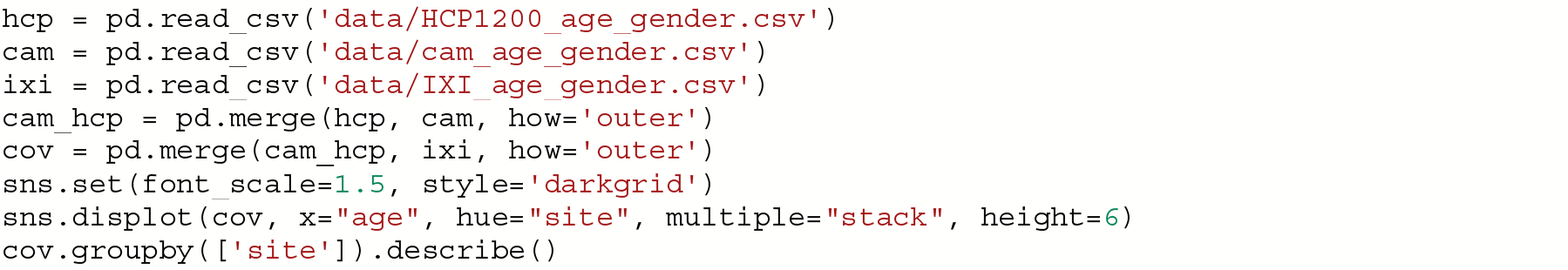

### Data Preparation: Prepare brain data

#### Timing: 10-15 minutes

3. Next, format and combine the MRI data using the following commands. The example data contains cortical thickness maps estimated by running recon-all from Freesurfer (version 6.0). The dimensionality of the data was reduced by using ROIs from the Desikan-Killiany atlas. Including the <u>Euler number</u> as a covariate is also recommended, as this is a proxy metric for data quality. The Euler number from each subject’s recon-all output folder was extracted into a text file and is merged into the cortical thickness data frame. The Euler number is site-specific, thus, to use the same exclusion threshold across sites it is important to center the site by subtracting the site median from all subjects at a site. Then take the square root and multiply by negative one and exclude any subjects with a square root above 10.

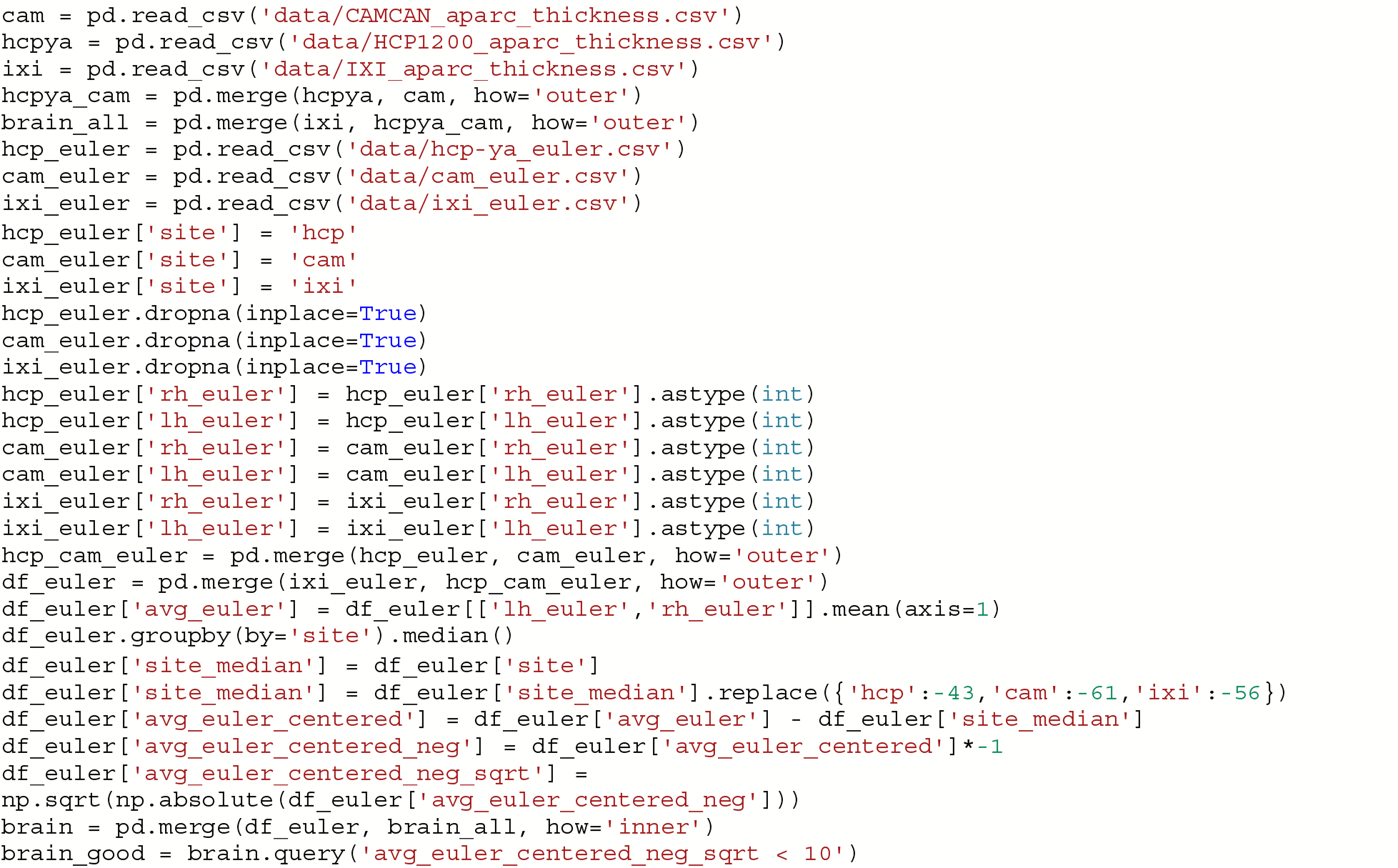

**CRITICAL STEP:** If possible, data should be visually inspected to verify that the data inclusion is not too strict or too lenient. Subjects above the Euler number threshold should be manually checked to verify and justify their exclusion due to poor data quality. This is just one approach for automated QC used by the developers of the PCNtoolkit.

Other approaches such as the <u>ENIGMA QC pipeline</u> or UK Biobank’s QC pipeline ^55^ are also viable options for automated QC.

### Data Preparation: Check that subjects (rows) align across covariate and brain dataframes

#### Timing: 3-5 minutes

4. The normative modeling function requires the covariate predictors and brain features to be in separate text files. However, it is important to first (inner) merge them together, using the following commands, to confirm that the same subjects are in each file and that the rows (representing subjects) align. This requires that both data frames have ‘subject_id’ as a column name. Once this is confirmed, exclude rows with NaN values and separate the brain features and covariate predictors into their own dataframes, using the commands below.

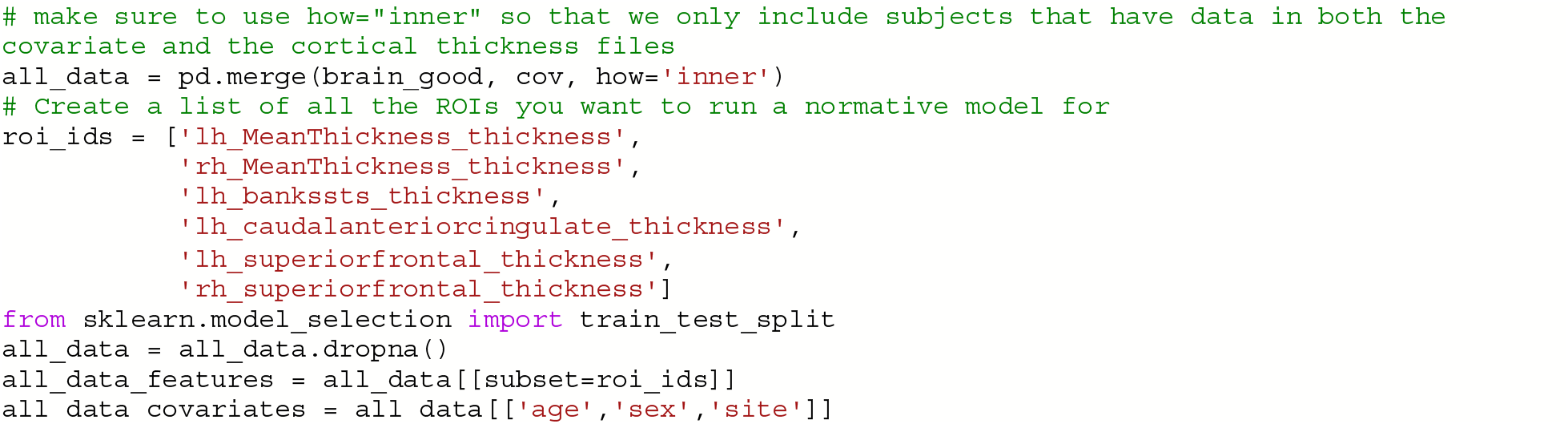

**CRITICAL STEP:** roi_ids is a variable that represents which brain areas will be modeled and can be used to select subsets of the data frame if you do not wish to run models for the whole brain.

### Data Preparation: Add variable to model site/scanner effects

#### Timing: 3-5 minutes

5. Currently, the different sites are coded in a single column (named ‘site’) and are represented as a string data type. However, the PCNtoolkit requires binary variables. Use the pandas package as follows to address this, which has a built-in function, pd.get_dummies, that takes in the string ‘site’ column and dummy encodes the site variable so that there is now a column for each site and the columns contain binary variables (0=not in this site, 1=present in this site).

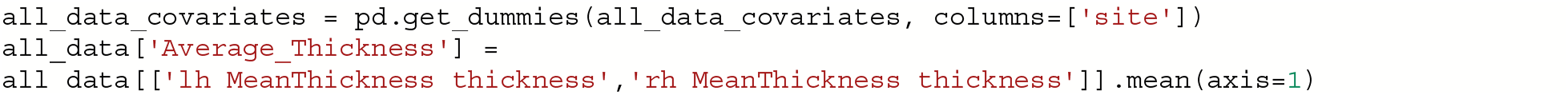

### Data Preparation: Train/Test split

#### Timing: 5-10 minutes

6. In this example, we use 80% of the data for training and 20% for testing. Please carefully read the experimental design section on train/test split considerations when using your own data in this step. Using a function from scikit-learn (train_test_split), stratify the train/test split using the site variable to make sure that the train/test sets both contain data from all sites, using the following commands. Next, confirm that your train and test arrays are the same size (rows), using the following commands. You do not need the same size columns (subjects) in the train and test arrays, but the rows represent the covariate and responses which should be the same across train and test arrays.

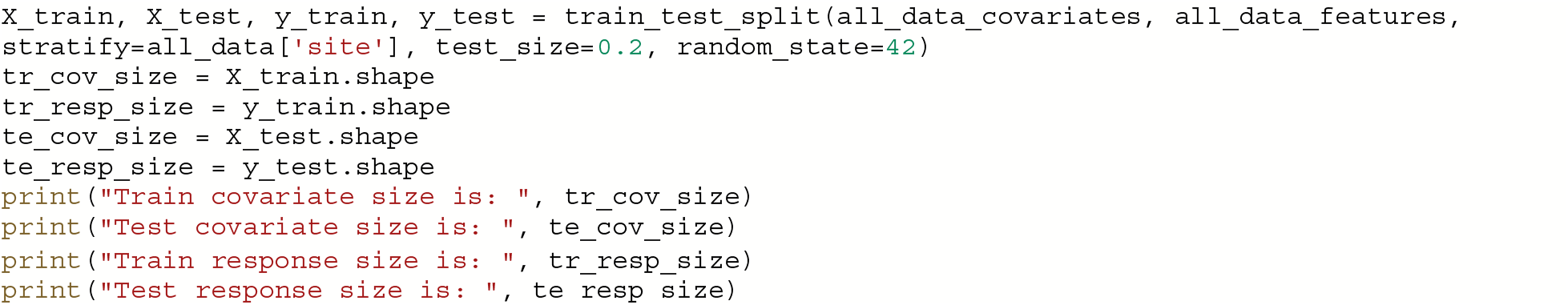

**CRITICAL STEP:** The model would not learn the site effects if all the data from one site was only in the test set. Therefore, we stratify the train/test split using the site variable.

7. When the data were split into train and test sets, the row index was not reset. This means that the row index in the train and test data frames still correspond to the full data frame (before splitting the data occurred). The test set row index informs which subjects belong to which site, and this information is needed to evaluate per site performance metrics. Resetting the row index of the train/test data frames fixes this issue. Then extract the site row indices to a list (one list per site) and create a list called site_names that is used to decide which sites to evaluate model performance for, as follows:

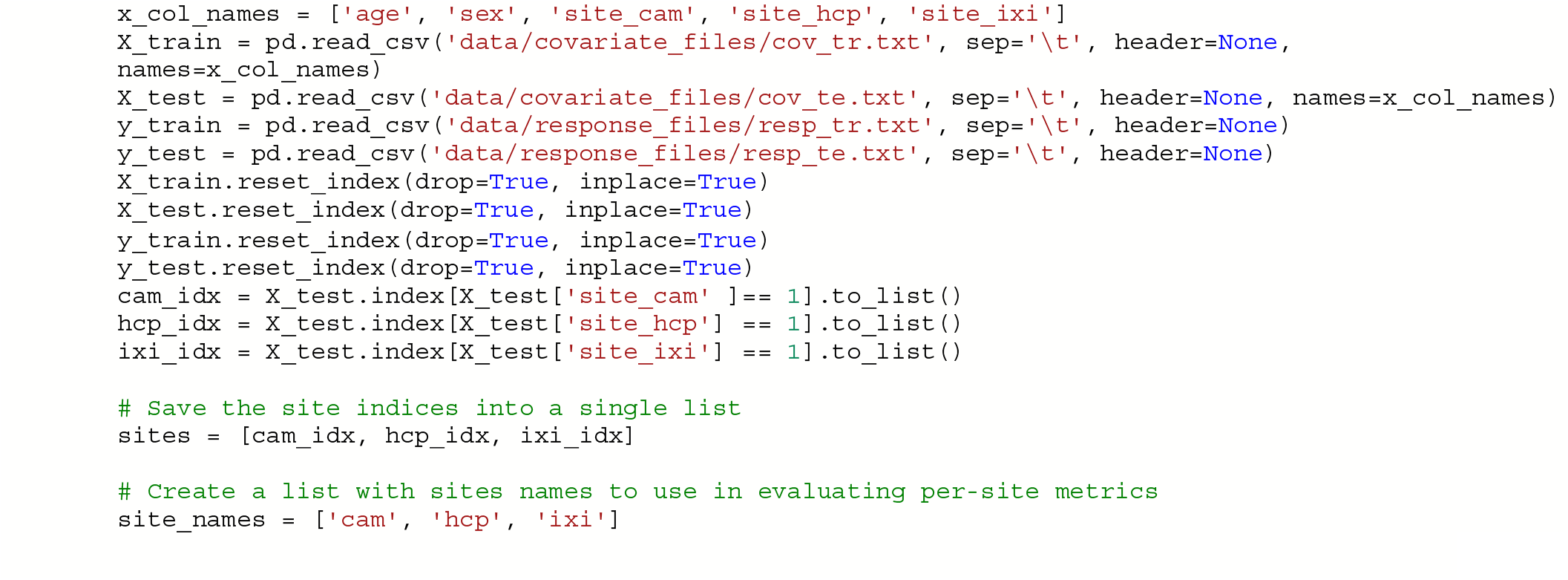

### Data Preparation: Setup output directories

#### Timing: 1-3 minutes

8. Save each brain region to its own text file (organized in separate directories) using the following commands, because for each response variable, **Y** (e.g., brain region) we fit a separate normative model.

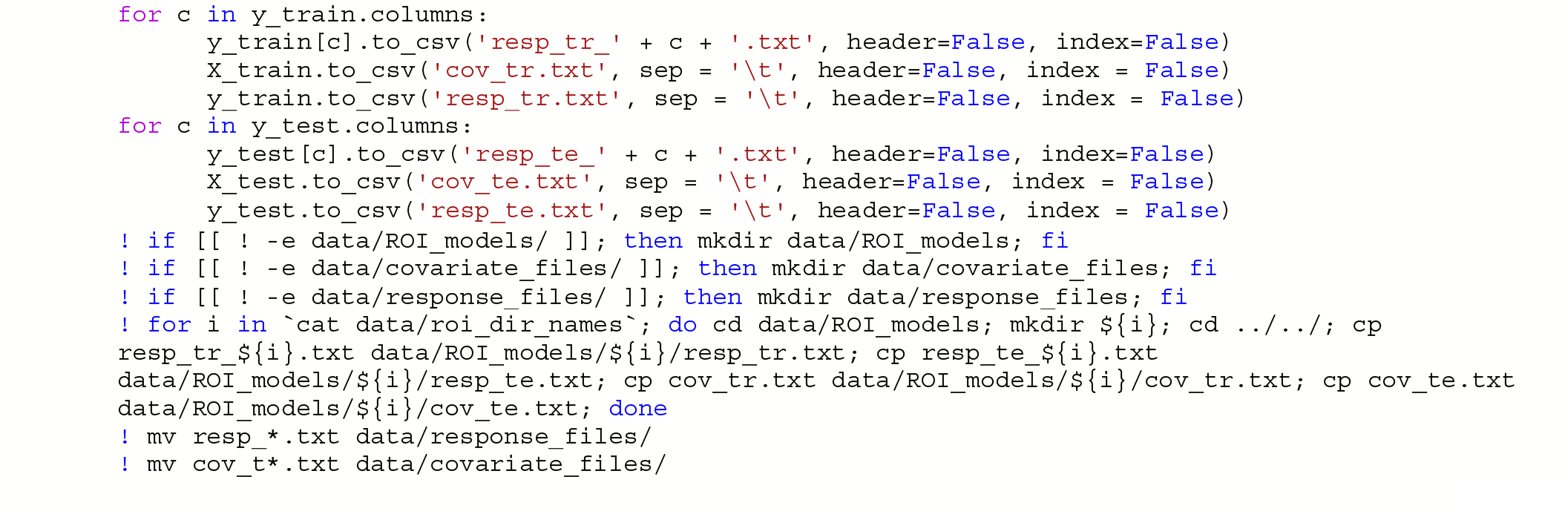

### Algorithm & Modeling: Basis expansion using B-splines

#### Timing: 1-3 minutes

9. Now, set up a B-spline basis set that allows us to perform nonlinear regression using a linear model, using the following commands. This basis is deliberately chosen to not to be too flexible so that it can only model relatively slowly varying trends. To increase the flexibility of the model you can change the parameterization (e.g., by adding knot points to the B-spline basis or increasing the order of the interpolating polynomial). Note that in the neuroimaging literature, it is more common to use a polynomial basis expansion for this. Piecewise polynomials like B-splines are superior to polynomial basis expansions because they do not introduce a global curvature. For further details on the use of B-splines see Fraza et al^25^.

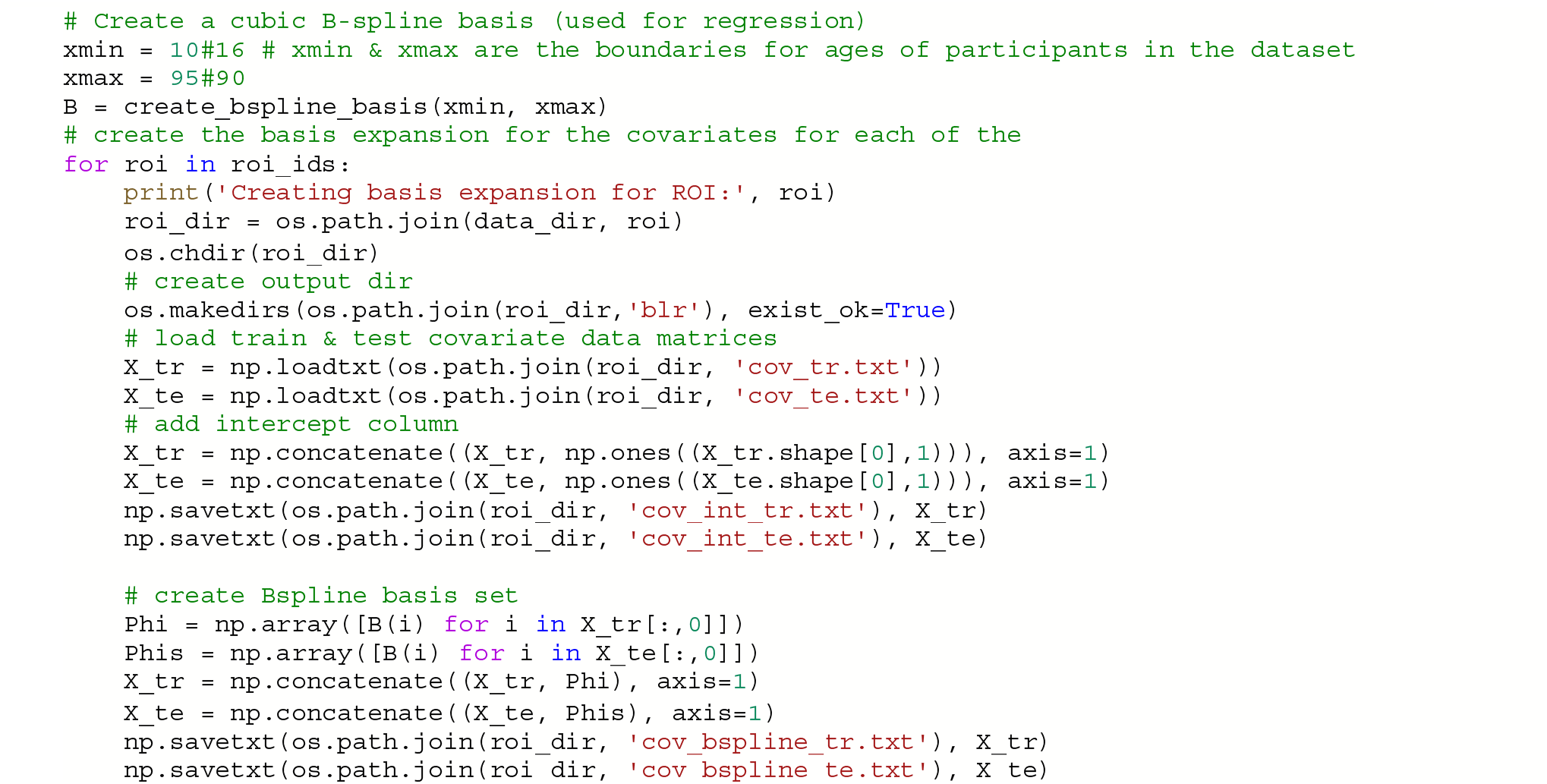

### Algorithm & Modeling - estimate normative model

#### Timing: 3-5 minutes per model (multiply by number of ROIs/models)

10. Set up a variable (data_dir) that specifies the path to the ROI directories that were created in Step 7. Initiate two empty pandas data frames where the evaluation metrics are the column names, as follows; one will be used for overall test set evaluation (blr_metrics) and one will be used for site-specific test set evaluation (blr_site_metrics). After the normative model has been estimated, these data frames will be saved as individual csv files.

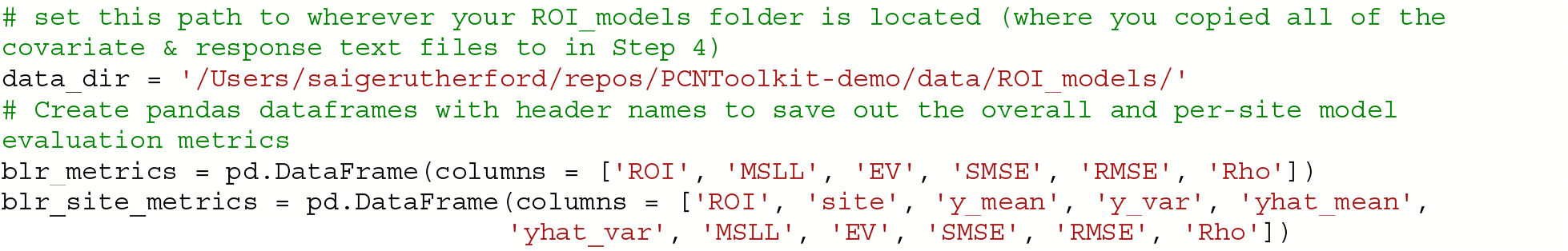

11. Estimate the normative models using a for loop to iterate over brain regions. The estimate function uses a few specific arguments that are worthy of commenting on:

- alg = ‘blr’: specifies we should use Bayesian Linear Regression. See Table 1 for other available algorithms.
- optimizer = ‘powell’: use Powell’s derivative-free optimization method (faster in this case than L-BFGS)
- savemodel = False: do not write out the final estimated model to disk
- saveoutput = False: return the outputs directly rather than writing them to disk
- standardize = False: Do not standardize the covariates or response variables

An important consideration is whether to re-scale or standardize the covariates or responses. Whilst this generally only has a minor effect on the final model accuracy, it has implications for the interpretation of models and how they are configured. If the covariates and responses are both standardized (standardize = True), the model will return standardized coefficients. If (as in this case) the response variables are not standardized (standardized = False), then the scaling both covariates and responses will be reflected in the estimated coefficients. Also, under the linear modeling approach employed here, if the coefficients are unstandardized and do not have a zero mean, it is necessary to add an intercept column to the design matrix (this is done above in step 9 (B-spline)).

**CRITICAL STEP:** This code fragment will loop through each region of interest in the roi_ids list (created in step 4) using Bayesian Linear Regression and evaluate the model on the independent test set. In principle, we could estimate the normative models on the whole data matrix at once (e.g., with the response variables stored in a n_subjects by n_brain_measures NumPy array or a text file instead of saved out into separate directories). However, running the models iteratively gives some extra flexibility in that it does not require that the included subjects are the same for each of the brain measures.

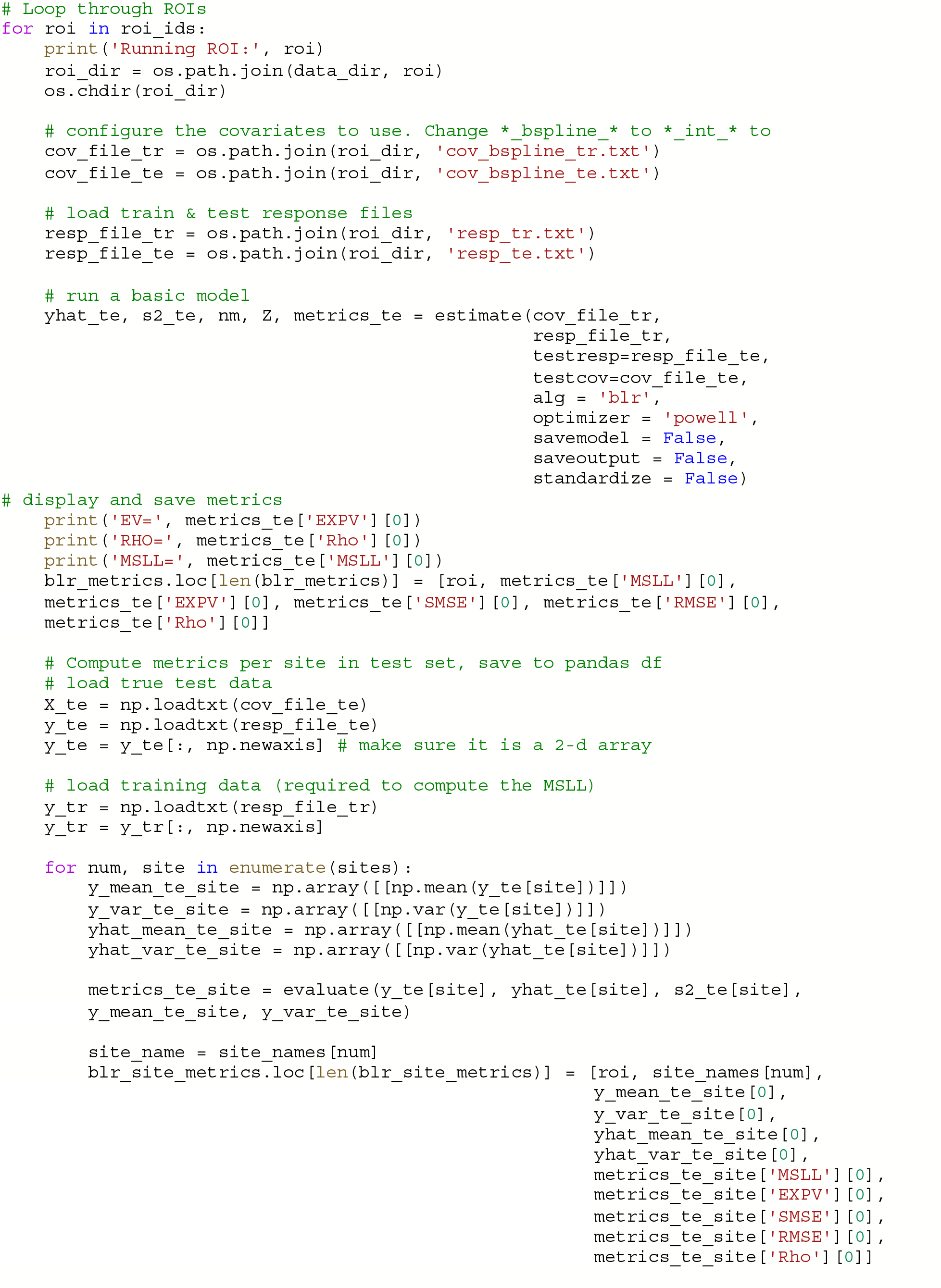

### Evaluation & Interpretation - evaluate normative model performance

#### Timing: 5-10 minutes

12. In step 11, when we looped over each region of interest in the roi_ids list (created in step 4) and evaluated the normative model on the independent test set, it also computed the evaluation metrics such as the explained variance, mean standardized log-loss and Pearson correlation between true and predicted test responses. The evaluation metrics were calculated for the full test set and calculated separately for each scanning site. The metrics were saved out to a csv file. In this step we load the evaluation metrics into a panads data frame and use the describe function to show the range, mean, and standard deviation of each of the evaluation metrics. Table 2 shows how to interpret the ranges/directions of good model fit.

**Table 2:**
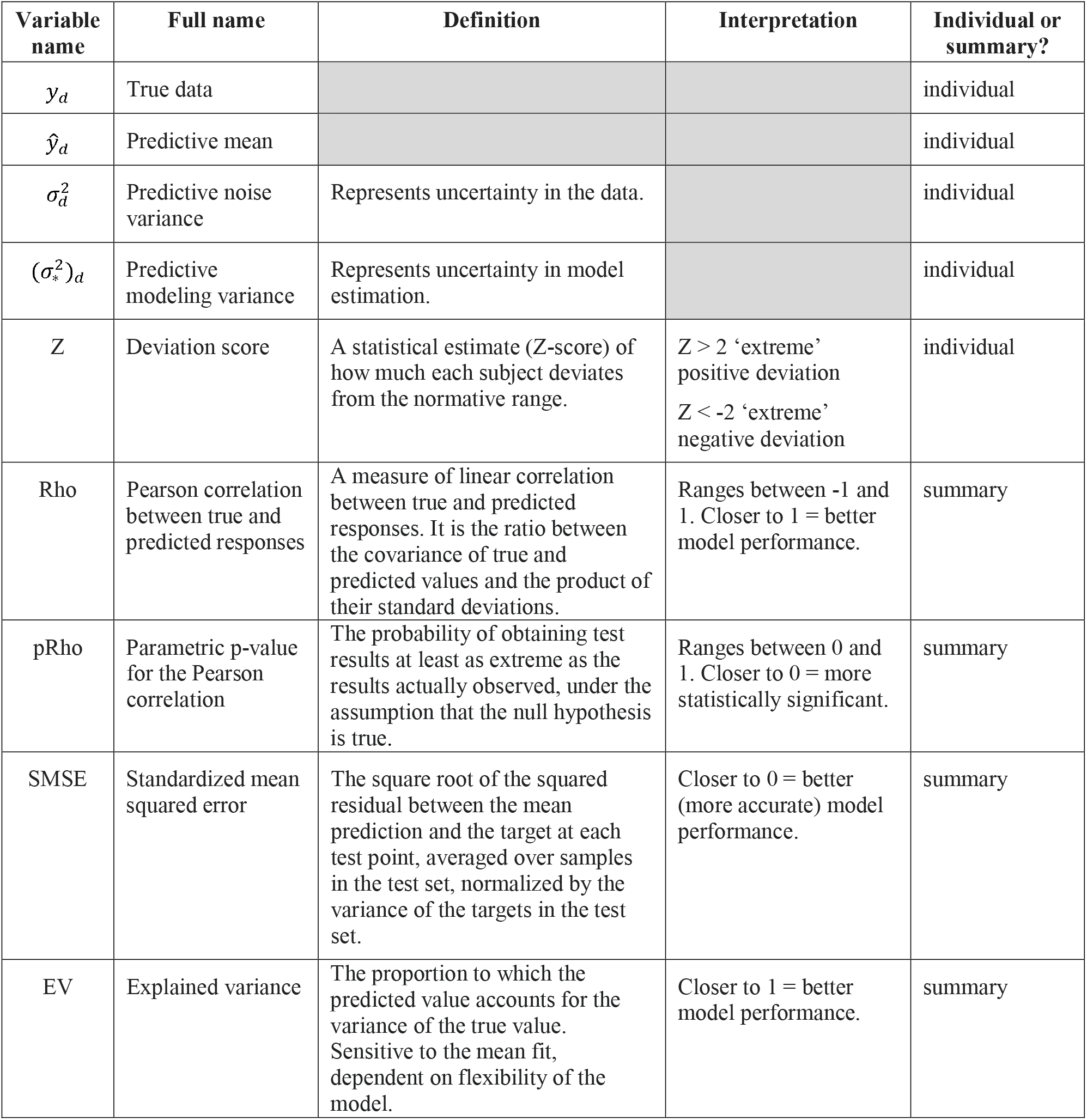
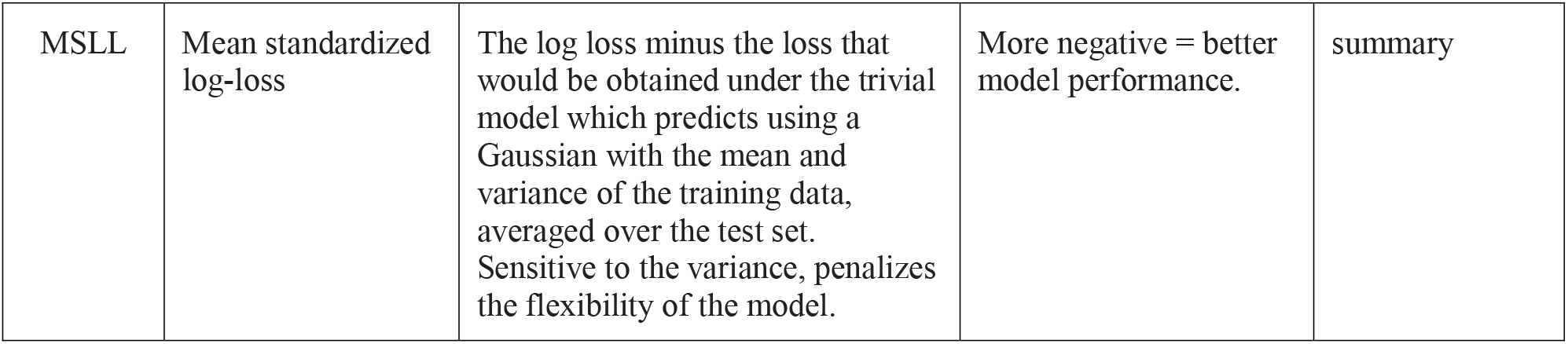
Normative Model Metrics. The **‘**Individual or summary?’ column refers to whether there is a value for every subject or if the metric is summarized across all subjects. For summary metrics, there is one value per brain region (model), and for individual metrics there are n_subjects x n_brain_regions values.

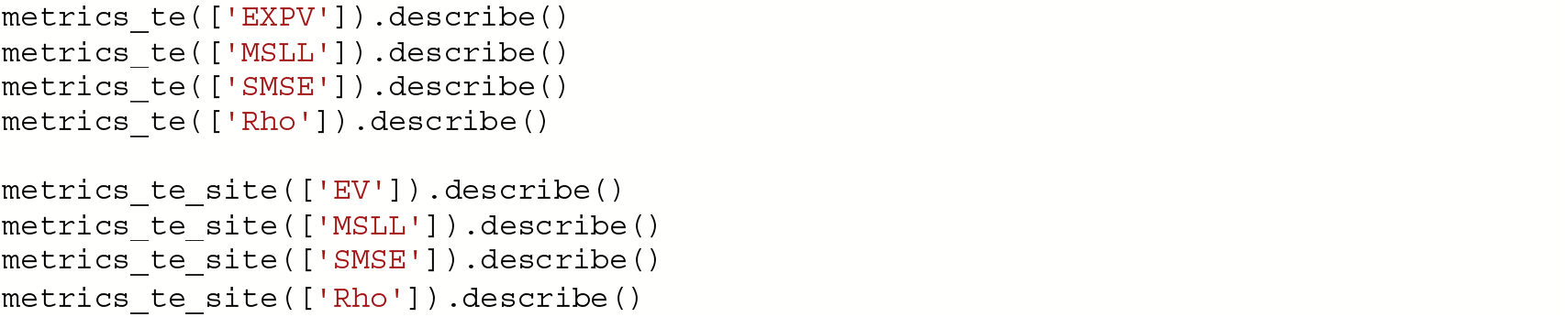

### Evaluation & Interpretation - visualize normative model outputs

#### Timing: 15-20 minutes

13. In this step we look at different ways of visualizing the evaluation metrics from step 12. There are typically many models fit across the different brain regions and it can be a lot of effort to keep track of the performance across all the brain regions. Data visualization will help to understand if there are any emerging patterns and find if there are any brain areas (or certain sites) where the model does not fit well. We summarize the deviation scores in the test set by counting how many subjects have an ‘extreme’ deviation (either positive or negative) and visualize the count of extreme negative and positive deviations by plotting them on a 3D brain plot. This step requires using a separate python notebook (https://github.com/predictive-clinical-neuroscience/PCNtoolkit-demo/blob/main/tutorials/BLR_protocol/visualizations.ipynb)

### Evaluation & Interpretation – post-hoc analysis ideas using normative modeling outputs

#### Timing: 1-2 hours

14. There are many interesting analyses that can be conducted using the outputs of normative modeling (deviation scores). An in-depth tutorial on each of these analyses is outside the scope of this protocol. However, on <u>GitHub</u>, we include code examples (python notebooks that can be run via Colab) of the following post-hoc analysis: (https://github.com/predictive-clinical-neuroscience/PCNtoolkit-demo/blob/main/tutorials/BLR_protocol/post_hoc_analysis.ipynb)

Using deviation scores as predictors in a regression and classification and comparing the performance to using the true data as predictors. Code for implementing several common predictive modeling frameworks (that are mentioned in comparison to other methods section) is provided. Deviation scores from normative modeling could be used as input features to any of these predictive modeling frameworks.
Using a pre-trained normative model and transferring it to a new, unseen data set.
Classical case-control testing (univariate t-tests) on deviation maps compared to univariate t-tests on the true data.

## Timing

The normative modeling portion of this protocol (including evaluation and visualization) can be completed in approximately 57-72 minutes. If using the additional code for post-hoc analysis of the normative modeling outputs, you would add approximately 1-2 hours to the estimated normative modeling time. These timing estimates are if using the Google Colab platform to run the code. If running this protocol on your own computer (where you need to install python and dependencies), this will add extra time to the protocol.

## Anticipated Results

There are multiple end products created from running a normative model analysis. First, the evaluation metrics for each model (brain region) are saved to a file. In this protocol, we saved the metrics to a CSV file format, however, in the pcn.estimate() function you could set the argument ‘binary = True’ which would save the metrics in pickle (.pkl) format. Pickle format is good to use if you are estimating many models in parallel on a large dataset, as it is faster because it avoids reading/writing intermediate text files. These metrics are further summarized into per site metrics to check model fit for each site included in the test set. The short and full names of the evaluation metrics and a brief interpretation guide is summarized in Table 2. The evaluation metrics can be visualized in numerous formats, histograms/density plots, scatter plots with fitted centiles, or brain-space visualizations. Several examples of these visualizations are shown in Figure 4 and code for creating these plots is shared on GitHub. Quality checking the normative model evaluation metrics should be done to ensure proper model estimation. If a model fits well to the data, the evaluation metrics should follow a Gaussian distribution. The model estimation (Procedure step 11) should properly handle confounding site effects, nevertheless, it is also a good idea to check per site metrics to make sure the model is fitting all sites equally well and that there are no obvious site outliers. In addition to the summary level evaluation metrics, there are also many individual metrics (one value per subject for each model/brain region). These individual-level outputs can be very helpful for interpretation because they precisely quantify the uncertainty of each individual predicted value at every location across the brain. If a given individual is identified as having an ‘extreme’ deviation, and there is low uncertainty you can be confident this is a biologically valid finding and not due to modeling errors. Vice versa, if there are extreme deviations and high levels of uncertainty, more caution should be given to interpretating these results and the deviations may be due to modeling errors rather than true biological variation. The uncertainty estimates are separated into two components (noise and modeling, described in Table 2) to help pinpoint the sources of uncertainty.

**Figure 4.**
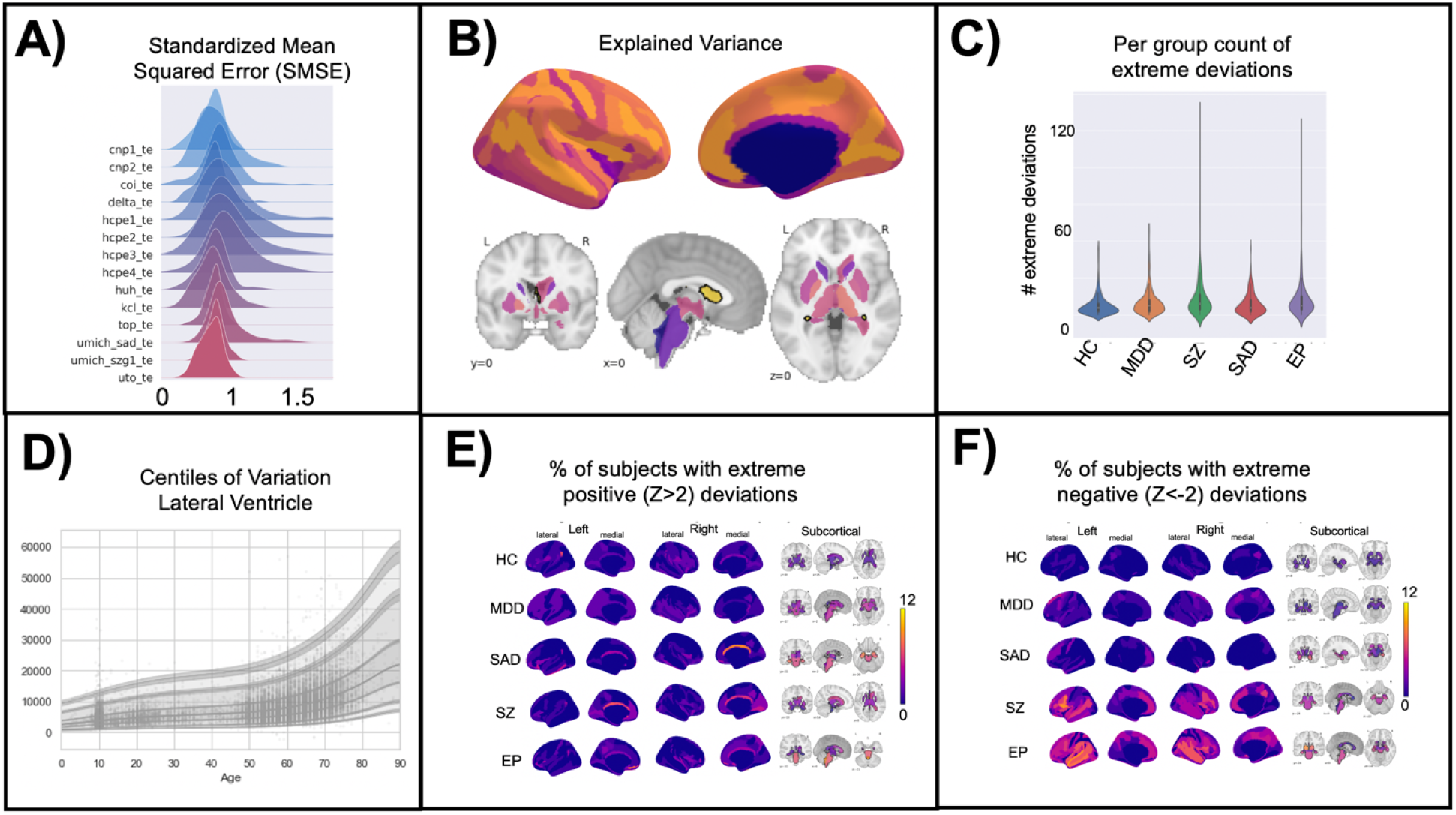
Visualization of Normative Model Evaluation Metrics. **A)** A ridge plot showing the distribution across all brain regions of the standardized mean squared error (SMSE), an evaluation metric that represents accuracy, visualized for each site in the test set. Visualizing for each test site can help identify if there are sites where the model is performing poorly. Ideally, the distribution will be Gaussian and should look similar across all sites. Small shifts in the mean across sites is to be expected and is acceptable. **B)** Explained variance is shown for cortical thickness of every brain region in the Destrieux parcellation) and volume of subcortical regions. Visualizing the evaluation metrics in brain space helps to identify patterns and see the big picture. **C)** The number of extreme deviations (both positive and negative) are counted for each individual in the test set, group ID is used to plot the distribution of the extreme deviation count for each group. A statistical test can be done on the count to determine if there is a significant difference between groups. Testing group differences in the count of deviations does not require there to be spatial overlap of the deviations within the group (i.e., this test can account for within-group heterogeneity of deviations). **D)** The normative trajectory for an example brain region (lateral ventricle) showing age (x-axis) versus the predicted volume (y-axis). The centiles of variation are shown by the lines and shaded confidence intervals. Each subject in the test set is plotted as a single point. **E-F)** Extreme deviations, separated into positive (**E**) and negative (**F**), are summarized for each group. For each brain region, the number of subjects with an extreme deviation in that region is counted, then divided by the group sample size, to show the percent of subjects with an extreme deviation. These visualizations demonstrate the benefit of normative modeling as there is within group heterogeneity that other methods (i.e., case-control group difference testing) are not equipped to handle. Abbreviations: HC = Controls, MDD=Major Depressive Disorder, SZ=Schizophrenia, SAD=Social Anxiety Disorder, EP=Early Psychosis.

A benefit of the PCNtoolkit software for normative modeling, that sets our approach apart from other normative modeling implementations^84^, is the fine-scale resolution allowed by the model. Other normative modeling work^84^ has focused on modeling gross features such as total brain volume or gray matter volume, which is not adequate for normative modeling applied to mental health conditions and neurodevelopmental disorders, where the effects are subtle and widespread (individuals within a patient group tend to deviate in different regions, see Figure 4E-F) across the cortex and subcortex and averaging over large brain areas usually overlooks these elusive psychiatric effects. This resolution also allows for a better mechanistic understanding because you can quantify the deviation and associated uncertainty for each individual with high spatial precision.

Reliability (the extent to which a measurement gives results that are very consistent) and validity (the degree to which a measurement measures what it is supposed to measure) are important constructs to keep in mind when interpreting results. In recent work, reliability of normative modeling in schizophrenia and bipolar disorder using structural MRI measures was established via replication^37^. Validity is arguably more challenging to assess but should be established by means of out of sample model fit. In other recent work, normative models were fit using a lifespan (age 3-100) big data sample (N=58,836) and carefully tested out-of-sample (variance explained, skewness, kurtosis, and standardized mean squared error) showing excellent model fit (12-68% variance explained) in an independent test set from a sample (and site) that was not included in the training set^85^. This work suggests validity, but this is an on-going evaluation and out of sample model fit must always be considered and reported.

## Troubleshooting

We re-iterate that there is additional documentation available online through <u>read the docs</u> including additional tutorials for other algorithm implementations (Gaussian Process Regression and Hierarchical Bayesian Regression), a glossary to clarify the jargon associated with the software, a reference guide with links to normative modeling publications, and a frequently asked questions page where many common errors (and their solutions) are discussed in detail. The problems encountered when troubleshooting a normative modeling analysis can fall into three categories: computing errors, data issues, and misunderstanding or misinterpreting the outputs.

## Computing errors

The computing errors might involve python or the computer hardware. Potential python errors may include installation of python or installation of the necessary packages and their dependencies. We recommend using Anaconda to install python 3.8 (required for this protocol) on your system, and the use of a virtual environment for the PCNtoolkit to ensure that the packages required for normative modeling do not interfere with other python versions and packages you may have installed on your system. In general, it is good to have a virtual environment setup for each project or analysis. If you are unfamiliar with setting up virtual environments, and run into issues with python, it is always an option to run the analysis in the cloud via Colab which eliminates the need to setup python on your own system. Hardware problems might include lack of memory to store the data or models running very slowly due to outdated hardware. These hardware errors do not have an easy solution, and we recommend using Google Colab to run normative modeling analysis if your personal computer or server is very slow or lacks the storage space.

## Data issues

Data issues that may be encountered are data missing not at random (see Experimental Design section regarding caution using data imputation), improperly coded data (i.e., strings instead of integers or floats, NaN values coded incorrectly), collinearity of columns in the covariate design matrix, or outlier data that does not make biological sense (i.e., negative cortical thickness values, negative age values). While these data errors can be incredibly frustrating to troubleshoot, they can typically be fixed by careful quality checking of the input data and removal of bad ROIs or subjects as needed.

### Interpretation confusion

Finally, an example of interpretation confusion may be poor model performance on a certain brain region or site. This can usually be addressed by returning to the input data for additional quality checking to confirm that the poor performance is not due to data quality issues. If there are no data quality issues, then it may be the reality that the model does not fit well some brain regions, and you may want to consider including additional covariates in the model to help explain more variance. Another interpretation confusion may arise when seeing negative explained variance values. When testing out of sample, the explained variance is not restricted to be positive, if it is negative this mean that the model fit is very poor (it is worse than an

intercept-only model).

## Code Availability Statement

All code is available on <u>GitHub</u> in the format of python notebooks that can be run in the cloud (for free) using <u>Google Colab</u>. We have also shared the GitHub repository on <u>Zenodo</u> to create a citable DOI for this software that also allows versions which are necessary as additional code and tutorials may be added over time^86^.

## Data Availability Statement

All data used in this protocol are available on <u>GitHub</u> and <u>Zenodo</u>^86^ in csv files. We also include a template csv file to help format user’s own data into the correct form for running the protocol using their own data set.

## Acknowledgements

This research was supported by grants from the European Research Council (ERC, grant “MENTALPRECISION” 10100118 and “BRAINMINT” 802998), the Wellcome Trust under an Innovator award (“BRAINCHART”, 215698/Z/19/Z) and a Strategic Award (098369/Z/12/Z), the Dutch Organisation for Scientific Research (VIDI grant 016.156.415). TW also gratefully acknowledges the Niels Stensen Fellowship as well as the European Union’s Horizon 2020 research and innovation programme under the Marie Sklodowska-Curie Grant agreement No. 895011.

## Author Contributions

Conceptualization: SR, SMK, TW, CF, MZ, RD, PB, AW, SV, HGR, CFB, AFM; Methodology: SR, SMK, TW, CF, MZ, RD, AFM; Data Curation: SR, AFM; Writing – Original Draft: SR; Writing – Reviewing and Editing: SR, SMK, TW, CF, MZ, RD, PB, AW, SV, HGR, CFB, AFM; Visualization: SR; Supervision: HR, CFB, AFM; Funding Acquisition: HR, CFB, AFM

